# EZH2 specifically regulates *ISL1* during embryonic urinary tract formation

**DOI:** 10.1101/2023.08.21.554100

**Authors:** Enrico Mingardo, Jeshurun C. Kalanithy, Gabriel Dworschak, Nina Ishorst, Öznur Yilmaz, Tobias Lindenberg, Ronja Hollstein, Tim Felger, Pierre-Olivier Angrand, Heiko Reutter, Benjamin Odermatt

## Abstract

*Isl1* has been described as an embryonic master control gene expressed in the pericloacal mesenchyme. Deletion of *Isl1* from the genital mesenchyme in mice leads to an ectopic urethral opening and epispadias-like phenotype. Using genome wide association methods, we identified *ISL1* as the key susceptibility gene for classic bladder exstrophy (CBE), comprising epispadias and exstrophy of the urinary bladder. The most significant marker (rs6874700) identified in our recent GWAS meta-analysis achieved a *p* value of 1.48 × 10^-24^ within the *ISL1* region. In silico analysis of rs6874700 and all other genome-wide significant markers in Linkage Disequilibrium (LD) with rs6874700 (D’ = 1.0; R^2^ > 0.90) revealed marker rs2303751 (*p* value 8.12 x 10^-20^) as the marker with the highest regulatory effect predicted. Here, we describe a novel 1.2 kb intragenic promoter residing between 6.2 and 7.4 kb downstream of the *ISL1* transcription starting site, which is located in the reverse DNA strand and harbors a binding side for EZH2 at the exact region of marker rs2303751. We show, that EZH2 silencing in HEK cells reduces *ISL1* expression. We show that *ezh2*^-/-^ ko zebrafish larvae display tissues specificity of ISL1 regulation with reduced expression of Isl1 in the pronephric region of zebrafish larvae. In addition, a shorter and malformed nephric duct is observed in *ezh2*^-/-^ ko zebrafish *Tg(wt1ß:eGFP)* reporter lines. Our study shows, that Ezh2 is a key regulator of *Isl1* during urinary tract formation and suggests tissue specific *ISL1* dysregulation as an underlying mechanism for CBE formation.

## Introduction

Using the *Isl1 (Islet1)*-Cre mouse line Suzuki et al. described *Isl1* as a master control gene expressed in the pericloacal mesenchyme (Suzuki et al., 2012). Ching et al. showed that deletion of *Isl1* from the genital mesenchyme in mice leads to hypoplasia of the genital tubercle and prepuce during early embryonic development, resulting in an ectopic urethral opening and epispadias-like phenotype (Ching et al., 2018). Meanwhile, using genome wide association methods, we identified *ISL1* as the key susceptibility gene for classic bladder exstrophy (CBE) (Draaken et al., 2015). CBE represents the most common defect form of the bladder exstrophy-epispadias complex (BEEC). About 98 % of all BEEC cases are classified as nonsyndromic respectively isolated (Beaman et al., 2021).The BEEC severity spectrum ranges from epispadias only, CBE to cloacal exstrophy and involves mainly the infraumbilical abdominal wall, the pelvis, all of the urinary tract, the genitals and in the severe cases spine and anus (Beaman et al., 2021). Our recent meta-analysis comprised seven independent discovery samples and identified eight genome-wide significant loci, seven of which were novel (Mingardo et al., 2022). The most significant marker (rs6874700) within the *ISL1* region achieved a *p* value of 1.48 × 10^-24^. Analysis of rs6874700 and all other genome-wide significant markers in LD with rs6874700 (D’ = 1.0; R^2^ > 0.90) revealed marker rs2303751 (*p* value 8.12 x 10^-20^) as the marker with the highest regulatory effect predicted (FORGEdb score 8 and RegulomeDB 2b; https://analysistools.cancer.gov/LDlink/?tab=ldproxy). ChIP-seq data indicate an EZH2 binding site at the genomic location of rs2303751 (doi:10.17989/ENCSR886KKK). EZH2 is the enzymatic catalytic subunit of PRC2 (Enhancer of Zeste 2 Polycomb Repressive Complex 2 Subunit) as well as a transcription factor (Kim et al., 2018). Our study now shows, that Ezh2 is a key regulator of *Isl1* expression during urinary tract formation. Overall, the findings of our study advance the understanding of normal urinary tract development and provide insights into the regulation of *ISL1*, the key susceptibility gene for CBE, and the underlying mechanisms of CBE formation.

## Results

### Luciferase assay identifies presence of a promoter in the reverse strand of the *ISL1* locus

We investigated the CBE associated marker rs2303751 using luciferase assays in HEK 293 cells covering a region of 2.854 base pairs (bp), located 5.028 to 7.882 bp downstream from the *ISL1* transcription starting site (TSS) (chr5:51388476-51391329, hg38) (Fig 1 B). We divided this region in three partially overlapping sequences that were cloned forward and flipped in pGL3-Basic plasmid for luciferase assays: (i) Fragment 1 ranged from 5.028 to 6.328 bp (1.300 bp) away from the *ISL1* TSS, (ii) Fragment 2 from 5.871 to 7.150 bp away from the *ISL1* TSS (1.279 bp), (iii) Fragment 3 from 6.610 to 7.882 bp away from of the *ISL1* TSS (1.272 bp) (Fig 1 B). Significant luciferase signal was observed only in the flipped Fragment 2 harboring rs2303751 (Fig 1 B and C). We then introduced the A>G minor allele variant of rs2303751 into Fragment 2 forward and flipped plasmids to test for luciferase activity. Here, neither the intensity nor the orientation of the promoter showed any differences compared to the effects of the major A allele (Fig 1 D).

**Figure 1.**
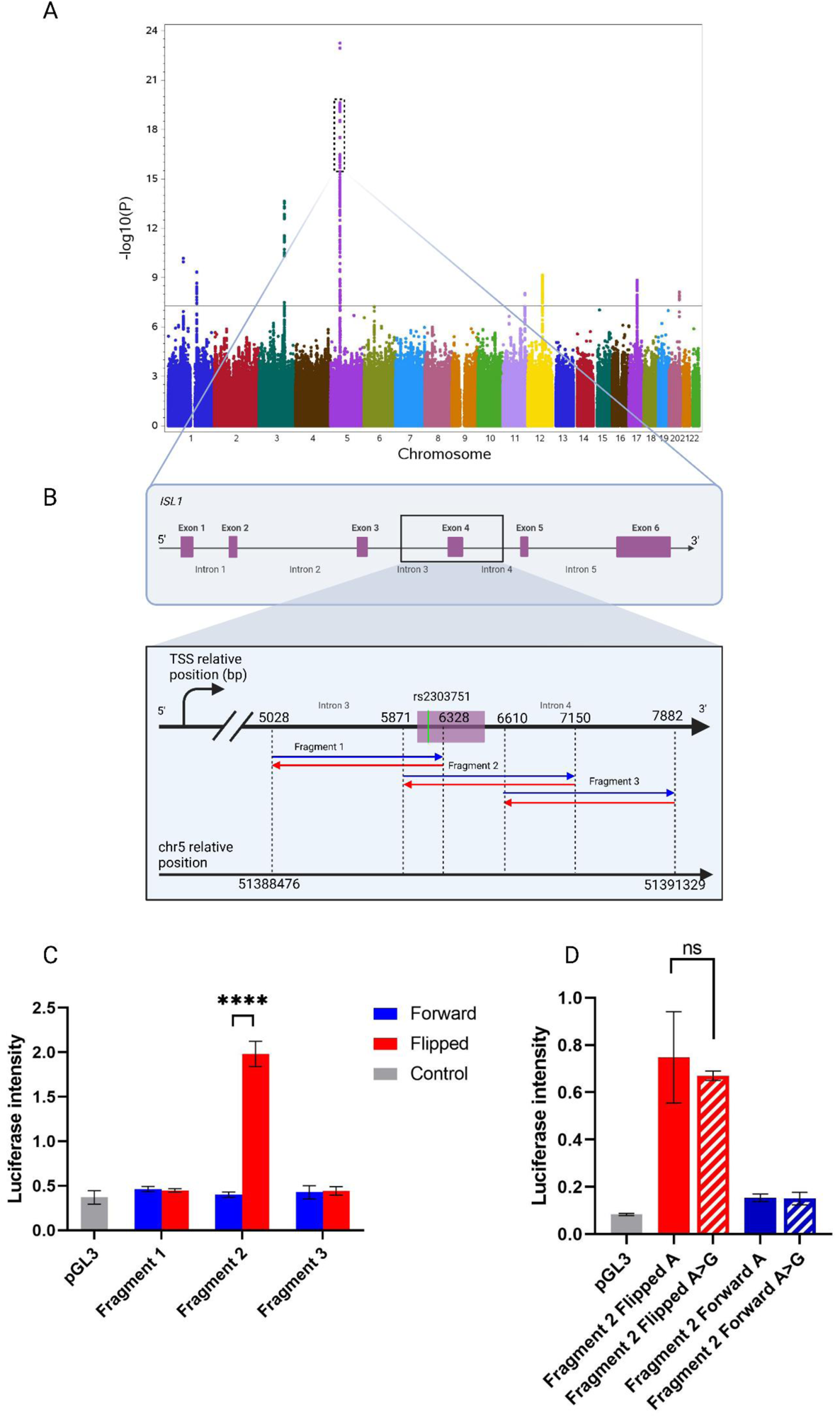
Luciferase sliding window approach identifies a promoter in the reverse strand of the rs2303751 harboring region. A) Manhattan plot of the CBE GWAS. The dotted box shows the region harboring *ISL1* (Mingardo et al., 2022) and is zoomed into B) Overview of the genomic region where the luciferase is performed referred to the *ISL1* gene. In detail, Fragment 1, Fragment 2 and Fragment 3 coordinates are shown relatively to the distance from the *ISL1* transcription starting site (TSS) (top) and also in chromosome 5 coordinates (hg38/lower). Blue and red arrows indicate the orientation of the luciferase-tested fragments. C) Luciferase assay of the forward (blue) and flipped (red) fragments relative to the empty control vector pGL3 (in gray); significance over control is observed only for the flipped Fragment 2. D) Fragment 2 luciferase assay with the rs2303751 variant A>G in both forward and flipped orientations displays no significant difference between major (A) and minor (G) allele.

### EZH2 protein binds to the GWAS associated CBE locus and regulates *ISL1* expression

ChIP qPCR in HEK 293 cells revealed presence of EZH2 binding to the *ISL1* genomic locus. In detail, we investigated 10 qPCR fragments equally distributed over a region that overlaps Fragment 2 from minus 1.6 kb upstream to 1.7 kb down-stream (Fig 2 A). We observed binding of EZH2 protein to the whole tested genomic region and found the highest fold change in binding for the two tested fragments of minus 0.5 kb and plus 0 kb distance of Fragment 2 (Fig 2 B). To further investigate the regulation of *ISL1* through EZH2 binding, we knocked down EZH2 in HEK 293 cells using siRNA (Fig 2 C) and tested ISL1 expression by qPCR. We observed a 1.5-fold decrease in *ISL1* expression following the EZH2 knockdown (Fig 2 D). To further check regulation of the Fragment 2 promoter, we overexpressed EZH2 using the pCMVHA expression plasmid (Fig. 2 E). We observed a significant doubling of luciferase activity in the flipped Fragment 2 version when overexpressing EZH2 but no effect on the empty controls nor on the Fragment 2 promoter in the forward orientation (Fig. 2 F).

**Figure 2.**
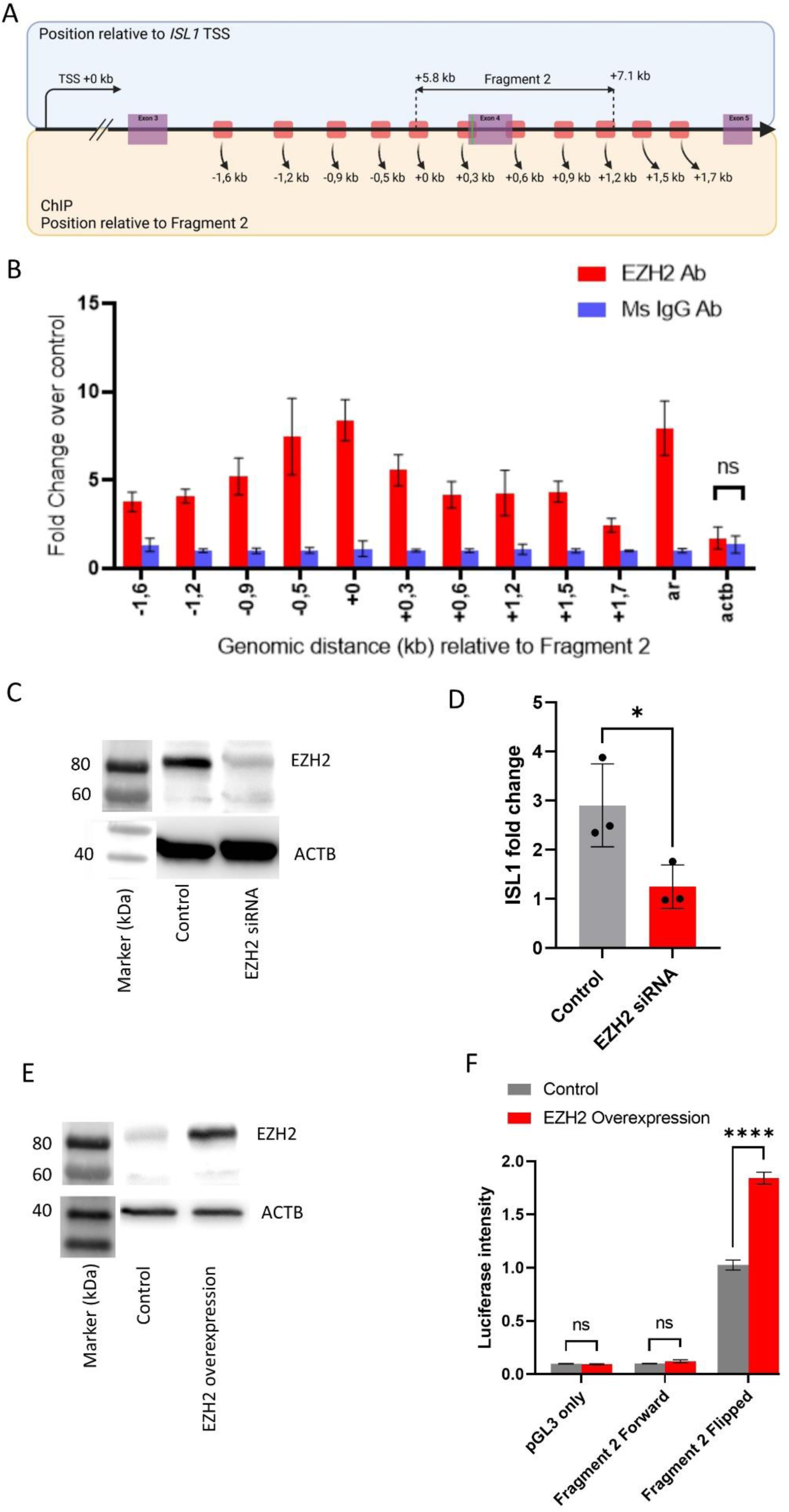
EZH2 enhance *ISL1* expression through binding on Fragment 2. A) schematic reppresentation of the region tested for ChIP-qPCR in *ISL1* (red segments) with distance relative to the 5’ of Fragment 2 in kb. B) ChIP-qPCR fold change representing EZH2 bound genomic DNA relative to mouse IgG bound DNA. Androgen receptor (ar) genomic DNA as positive control (Kim et al., 2018) and actin b as negative control. All fold change except for actin b are significant over IgG control. C) Western blot against EZH2 shows a succefull siRNA knockdown of EZH2 compared to control. D) *ISL1* qPCR with EZH2 siRNA dispalys a reduced *ISL1* signal. E) Western blot on EZH2 with control vector and EZH2 overexpression plasmid confirms the efficency of the overexpression. F) Luciferase assay with EZH2 overexpression (red bars) and control (gray bars). As before, vector (pGL3) and Fragment 2 forward show only little expression in both cases while EZH2 overexpression enchances the reporter activitiy of the plasmid with flipped Fragment 2 significantly.

### Whole mount *in situ* hybridization (WISH) and immuno histochemistry on *ezh2^ul2-/-^* zebrafish larvae (zfl) displays tissue specific regulation of *isl1* and abnormal nephron development

To further investigate *isl1* regulation via Ezh2 in a vertebrate model, we performed mRNA *in situ* hybridization against *isl1* transcript in wt, *ezh2^ul2+/-^* (*ezh2^+/-^*) and *ezh2^ul2-/-^* (*ezh2^-/-^*, knock out (KO) (Dupret et al., 2017)) zebrafish larvae (zfl) at 56 hpf to study *isl1* regulation of the pronephric region (Zhang et al., 2017). We observed a decreased *isl1* expression, which locates specifically to the nephron region of the *ezh2^u/2-/-^* zfl only (Fig 3 A). There is no such visible strong decrease of *isl1* expression in the brain and spinal cord of the *ezh2^ul2-/-^* zfl. qRT-PCR of the trunk from head-chopped embryos, which still included the pronephric region, confirmed a significant decreased in *isl1* transcript signal in the knockout (KO) zfl (Fig 4 B). Whereas, the same qRT-PCR protocol using whole larvae did not display any significant differences in *isl1* expression between WT and *ezh2^ul2-/-^* zfl (Fig 3 C), which confirms our WISH finding of mainly pronephric regulation of *isl1*, possibly mediated via Ezh2. To confirm the presence of Isl1 protein within the pronephric region, we performed anti Isl1 immuno histochemistry on paraffin sections of *Tg(wt1b:GFP) ezh2^ul2+/+^* and *ezh2^ul2-/-^* double transgenic zfl at 56 hpf. Here, we observed a strong reduction of Isl1 protein specifically in cells of the glomeruli and nephric duct of the *ezh2^ul2-/-^* zfl (Fig 3 D). In *Tg(wt1b:GFP): ezh2^ul^* double transgenic zfl, we investigated the Ezh2 KO effect on the development of the pronephros. Here, we observed in all KO larvae an irregular and malformed nephric duct from 3 dpf onwards (Fig 3 E).

**Figure 3.**
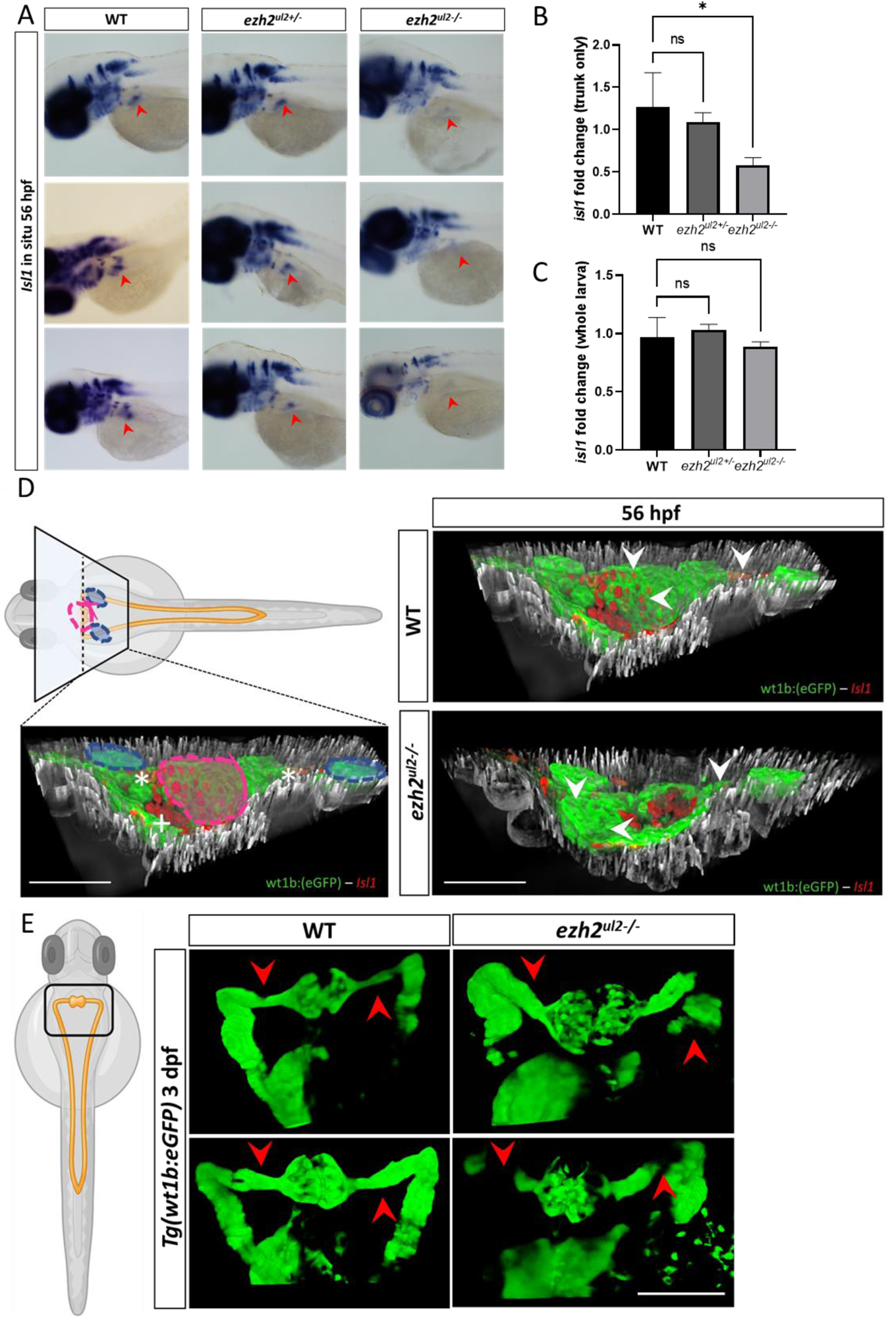
*Ezh2* mediates *isl1* regulation with tissue specificity on the nephric region and causes defective nephric duct development. A) *isl1* in situ hybridization in WT, Ezh2 +/- and KO larvae at 56 hpf. Red arrows indiacate the expression of *isl1* in the nephric region that shows clear staining in the WT and Ezh2+/- lines and a strong reduction in the KO zfl. *Isl1* expression results almost not alterated in the brain and spinal chord of all genotypes. B) *isl1* qPCR in head-chopped embryo at 56 hpf shows a significantly reduced signal in the Ezh2 KO line. C) *isl1* qPCR of whole zfl at 56 hpf shows no significant reduction of *isl1* expression. D) Isl1 immuno histochemistry (red cells) in *Tg(wt1b:eGFP)* line (in green) and double transgenic *Tg(wt1b:eGFP)* – Ezh2 KO line. Left pannel indicates the location of the transversal paraffin section in reference of the whole zfl and nephric region. Blue circles indicate the sagittal nephric ducts; purple circle indicates the glomeruli region, white asterics the sagittal nephric ducts and the white plus the pancreas. Right pannels show the 3D co-localization of *Isl1* protein (red) on the glomeruli and nephric ducts (green) in the WT (top) and *Ezh2* KO (lower) 56 hpf zfl. A clear absence of *Isl1* signal locates to the glomeruli and nephric ducts of the *Ezh2* KO line compared to WT. E) Nephric ducts of the *Ezh2* KO larvae display developmental defects and malformation at 3 dpf. Red arrows indicate the correct protrusion of the nephric duct in the WT and absence of GFP signaltogeather with dilated nephric ducts in the Ezh2 KO.

**Figure 4.**
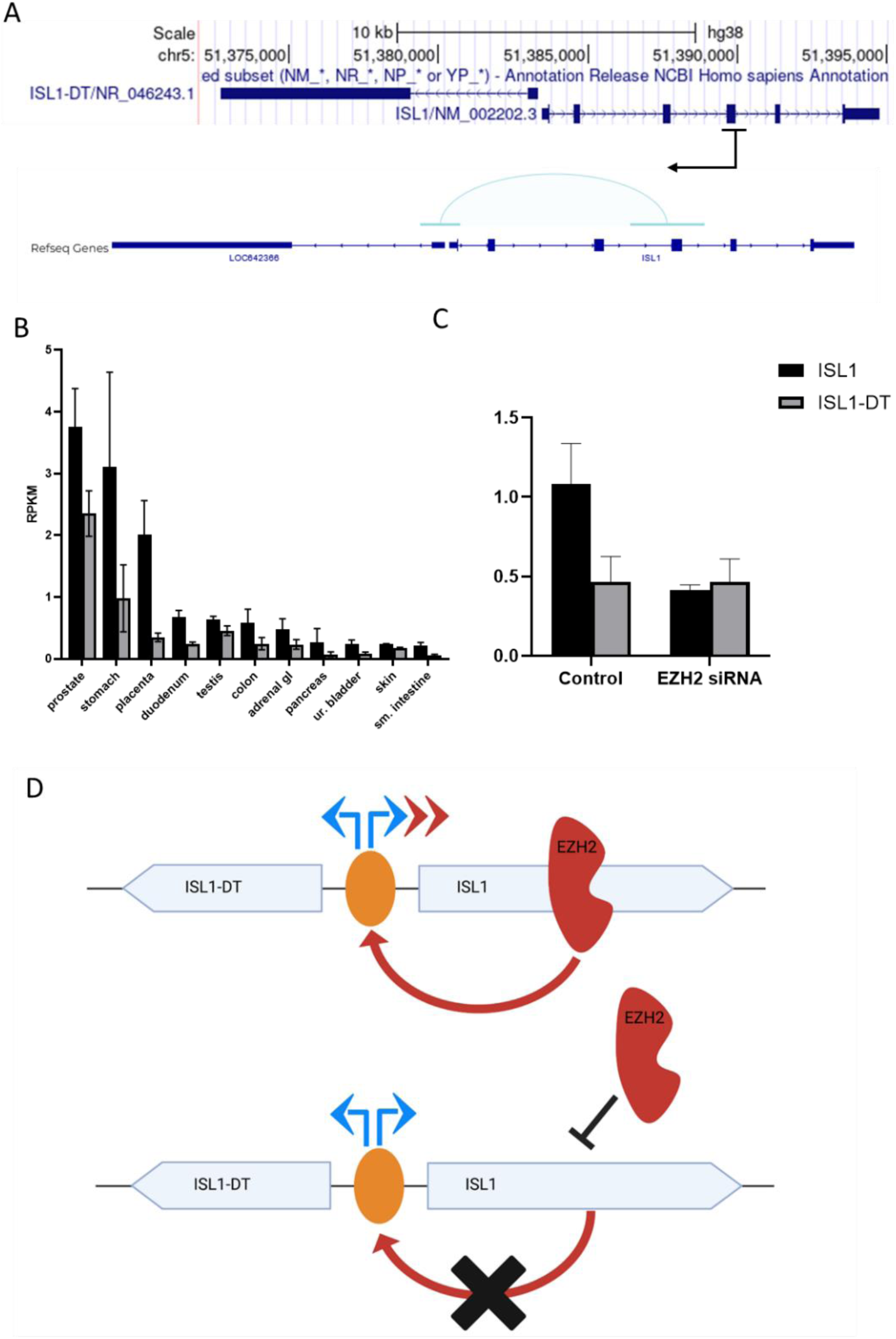
Fragment 2 promoter regulates specifically *ISL1* expression through EZH2 and disturbs the normal basal expression of *ISL1-DT/ISL1* cluster. A) Upper panel shows the genomic overview of *ISL1* location with 115 bp upstream the reverse orientated *ISL1-DT*. Black stripe indicates the position of Fragment 2 and the arrow its active orientation. Lower panel displays chromatin interaction from sci-ATAC-seq3 for *ISL1* in the fetal ureteric bud cells (Domcke et al., 2020). Light blue bridge indicates the interaction of the genomic locus where Fragment 2 resides with the shared promoter region of *ISL1* and *ISL1-DT* (light blue bridge). B) Human adult RNA-seq shows the expression pattern of *ISL1* and *ISL1-DT* in different tissues. These genes are generally expressed with higher counts for *ISL1* and lower for *ISL1-DT*. C) qPCR of *ISL1* and *ISL1-DT* with control and EZH2 siRNA shows a reduced signal of *ISL1* but unchanged *ISL1-DT* expression. D) Proposed molecular mechanism for the expression pattern of *ISL1* and *ISL1-DT* mediated by EZH2. Top panel shows physiological condition, EZH2 binds to the Fragment 2 promoter and specifically enhances *ISL1* expression (red arrows). In pathological condition, EZH2 does not bind to the Fragment 2 and the expression of *ISL1* is reduced, but not the expression of *ISL1-DT*.

### Fragment 2 promoter is specific for *ISL1* expression and dysregulates the ratio of *ISL1-*DT/*ISL1* expression

To further investigate the above described tissue specific effect and to further explore the mechanisms of *Isl1* regulation, we aimed to understand why Fragment 2 promoter is only active in its flipped orientation. The genomic region containing Fragment 2 has been shown to interact with the 115-bp region separating *ISL1* from *ISL1-DT* (*ISL1* Divergent Transcript) mediated by EZH2. *ISL1-DT* is an RNA gene affiliated with the long non-coding RNA (lncRNA) class. Fragment 2 in the reverse strand of *ISL1* orientation has regulatory effects. Interestingly, *ISL1-DT* is oriented in the reverse strand of *ISL1* as well. Because of this, we investigated whether the newly identified promotor has regulatory impact on *ISL1-DT* (Fig. 4 A). Previously, we found *Isl1-dt* to be significantly downregulated during embryonic urinary bladder development of mice (Mingardo et al., 2022). To investigate the expression pattern of *ISL1* and *ISL1-DT* we analyzed RNA-seq deposited data provided by the NCBI project “HPA RNA-seq normal tissues” (BioProject: <u>PRJEB4337</u>) (Fagerberg et al., 2014). Here we found that *ISL1* and *ISL1-DT* are expressed with the same basal expression pattern in the same analyzed tissues but *ISL1* always shows higher reads per kilo base of transcript per million mapped (RPKM) than *ISL1-DT* (Figure 4 B). This mechanism identifies those two genes as divergent lncRNA/mRNA transcripts (Sigova et al., 2013). To investigate whether the EZH2 mediated Fragment 2 promoter regulates both *ISL1-DT* and *ISL1* in a joined manner, we performed qPCR in HEK293 cells with and without EZH2 siRNA mediated knockdown. In the scrambled control we observed that both, *ISL1-DT* and *ISL1*, show the previously described basal expression of the analyzed “HPA RNA-seq normal tissue”, with *ISL1* showing approximately twice as high expression than *ISL1-DT*. While EZH2 knockdown does not change *ISL1-DT* expression level, the expression level of *ISL1* is decreased (Figure 4 C and Figure 2 D), indicating, that the EZH2 mediated Fragment 2 promoter does not regulate *ISL1-DT* but only *ISL1* expression (Figure 4 D).

## Discussion

Classic bladder exstrophy CBE represents the most common defect form of the bladder exstrophy-epispadias complex (BEEC). Embryonic formation is assumed to occur between the first four to six weeks of gestation. Recently, we identified *ISL1* as the key susceptibility gene for CBE (Draaken et al., 2015; Mingardo et al., 2022). Accordingly, the role of *ISL1* in urinary tract formation has been demonstrated by induced KO in mice leading to hypoplasia of the genital tubercle and prepuce during early embryonic development resembling an epispadias-like phenotype (Ching et al., 2018) (Su et al., 2019). However, no deleterious genetic variant within the *ISL1* gene nor copy number variations affecting the *ISL1* gene have been found in CBE individuals to date (Arkani et al., 2018) (Nordenskjöld et al., 2023). Hence, regulatory effects may be involved in CBE formation rather than sequence or copy number alterations of *ISL1*. As mentioned earlier, the marker rs2303751 was the genome-wide significant marker that was in LD (D’ = 1.0; R2 > 0.90) with the top marker rs6874700 (Supplementary Figure 1 A), which was predicted to have the highest regulatory effect within the *ISL1* gene and the surrounding region. The regulatory effect of marker rs2303751 was predicted to be much higher than that of the top marker rs6874700 itself (FORGEdb score 8 and RegulomeDB 2b; https://analysistools.cancer.gov/LDlink/?tab=ldproxy). ChIP-seq data suggests rs2303751 to define a binding site for EZH2 (Encode ID: <u>ENCSR000ATA</u> and <u>ENCSR850KIP</u>). EZH2 is well known as the enzymatic catalytic subunit of the PRC2 complex (Polycomb Repressive Complex Subunit 2) and recently it was described to act as a transcription factor (Kim et al., 2018). In this study we identified a promoter (here called Fragment 2) that is located in the rs2303751 defined region, which is active only in the reverse strand of the *ISL1* gene. In HEK 293 this region presents the histone marker H3K4me3 (Supplementary Figure 1 B; Encode ID: ENCSR000DTU) known to be associated with promoter-like loci (Bernstein et al., 2006) and we confirmed by ChIP-qPCR in HEK 293 cells, that EZH2 protein binds and occupies the rs2303751 defined region. We demonstrate that EZH2 knockdown in HEK 293 and KO in zf larvae downregulates *ISL1* expression. Remarkably, overexpression of EZH2 in HEK 293 does not increase ISL1 expression (Figure 2 F and Supplementary Figure 2) but it enhances the luciferase activity of the Fragment 2 (Figure 2 F). For this reason, we believe that the natural amount of intrinsic EZH2 is already saturating Fragment 2, and that an overexpression of EZH2 has no further impact on the genomic region of *ISL1*.

Since we found the Fragment 2 promoter active only in the reverse strand of *ISL1*, we investigated if its regulation through EZH2 could influence *ISL1-DT*, a long lncRNA that resides in the reverse strand 115 bp upstream to *ISL1*. We found that the two genes are always expressed with a constant basal pattern ratio showing a higher RPKM for *ISL1* and lower for *ISL1-DT*, remarkably there are no tissues where only the first or the latter are singularly expressed, suggesting that the 2 genes share the same promoter and are expressed simultaneously acting as described for divergent transcripts lncRNA/mRNA (Seila et al., 2008). We showed this pattern to be present in HEK293 cells as well (Figure 3 B), and that the EZH2 knockdown affects only *ISL1* expression while *ISL-DT* remains unchanged. This observation suggests a specific regulatory effect of Fragment 2 via EZH2 for *ISL1* only, suggesting this region to enhance specifically the *ISL1* gene expression and not *ISL1-DT*. The role of the divergent binomial expression of lincRNA/mRNA has been previously reported to play a role in endoderm specification of stem cells (Sigova et al., 2013). The urinary bladder develops through mesenchymal-epithelial interactions between the endoderm of the urogenital sinus and mesodermal mesenchyme (Liaw et al., 2018). Hence, dysregulation of the basal transcription between *ISL1* and *ISL1-DT* during development could contribute to a defect in the endoderm derived bladder tissue. To further explore a possible male-development caused by the EZH2-mediated *ISL1* downregulation, we investigated *ISL1* expression in *ezh2* zebrafish KO larvae *ezh2^ul2-/-^*. As previously reported by our group, *isl1* is expressed in the pronephros in 56 hpf zfl (Zhang et al., 2017). Therefore, we performed an *in situ* against *isl1* in ezh2^ul2-/-^ line and observed a reduced *isl1* signal that locates specifically to the pronephric region. This tissue specific reduction is additionally confirmed by q-rtPCR. Tissue specificity of *isl1* regulation through Ezh2 is further shown in our anti Isl1 immunohistochemistry analysis in the double transgenic zf line *Tg(wt1ß:eGFP):ezh2^ul2-/-^* where a reduction of Isl1 protein is located specifically to the pronephros but e.g. not in the spinal cord. In addition, *Tg(wt1ß:eGFP):ezh2^ul2-/-^* zfl display tissue anomalies and developmental defects within the nephric region starting at 3 dpf. These findings in the zf model suggest that the regulation of *isl1* via Ezh2 is tissue specific. We think that this tissue specificity is representative for vertebrates, as we find a similar expression in human derived HEK cells. Remarkably, when compared to human, neither the zf genome nor the mouse present a lnc-divergent RNA to *ISL1*, but still the *ISL1*-EZH2-mediated regulation shows conservation between zf and human as such. This suggests that EZH2 mediated regulation of *ISL1* represents an ancient regulatory mechanism that has been further transformed to and refined in human. Since non-coding divergent transcripts are known to regulate related nearby genes during embryonic tissue differentiation (Lepoivre et al., 2013), we assume that in human *ISL1-DT* has synergistic functions to *ISL1* in regulating differentiation of organs and tissues like prostate, stomach, placenta (as shown in Figure 3 B) and that specific EZH2-mediated dysregulation of *ISL1* in the *ISL1/ISL1-DT* cassette could contribute to developmental defect of the human urinary tract.

### Summing up

Our findings suggest that specific tissue dependent dysregulation of *ISL1* might be involved in CBE formation during early urinary bladder development and that this mechanism employs tissue specific regulation of *ISL1* through binding of EZH2 on the *ISL1* intrinsic Fragment 2. Hence, when EZH2 binding is impaired to this region, *ISL1* expression gets dysregulated interfering with the physiological development of the urinary tract. How EZH2 binding might be impaired, and how exactly the disturbance of the *ISL1* to *ISL1-DT* ratio may affect this developmental process should be the subject of future research.

## Materials and Methods

### Cell culture

Human embryonic kidney cells (HEK 293) were cultured in Dulbecco’s Modified Eagle Medium (DMEM, ThermoFisher) containing 10 % fetal bovine serum (FBS, PAN biotech), at 37°C and 5 % CO_2_ without antibiotics.

### Zebrafish husbandry

Adult zebrafish were maintained at 28°C with a light/dark cycle of 14/10 h under provided husbandry license of the Bonn medical faculty core facility (§ 11, City of Bonn, Germany). Embryos were gained by natural spawning and kept until 5 days post fertilization (dpf) at 28°C in an incubator in 0.3× Danieau’s buffer (1× Danieau’s buffer: 58 mM NaCl, 0.7 mM KCl, 0.4 mM MgSO4, 0.6 mM Ca(NO3)2, 5 mM HEPES, pH 7.2). Until 24 hpf, 0.3× Danieau’s buffer was supplemented with 0.00001 % methylene blue solution. From 24 hpf, when used for imaging, 0.3 × Danieau’s buffer was supplemented with 0.003 % phenylthiourea (PTU) to prevent pigmentation (Danieau/PTU solution). All experiments were done according to institutional and national law (https://www.nc3rs.orlineg.uk/arrive-guidelines). Transgenic fish lines *Tg(wt1b:eGFP)*, *ezh2^ul2+/--^* TALEN derived line (Dupret et al., 2017) and *Tg(wt1b:eGFP)-ezh2^+/-^* were used in this study.

### Dual luciferase assays

Genomic DNA was extracted from HEK 293 cells with DNeasy Blood & Tissue Kit (Qiagen, Cat. No. / ID:69504) and PCR amplified with primers specific for the investigated *ISL1* region (primers list in supplementary information). Fragments were inserted in reporter vector pGL3-Basic Vector (Promega, Cat. No. E1751) upon XhoI and MluI-HF enzymes digestion (New England BioLabs, respectively Cat. No. R0146L and R3198L). QuikChange Lightning Site-Directed Mutagenesis Kit (Agilent, Cat. No. 210518) was used to insert the variant of interest rs2303751 in the plasmids. In total 8 reporter vectors were cloned according to standard procedures: pGL3-Fragment1 (forward and flipped; chr5:51388476-51389775), pGL3-Fragment2 (forward and flipped; chr5:51389319-51390597), pGL3-Fragment2 (forward-rs2303751 and flipped-rs2303751), pGL3-Fragment3 (forward and flipped; chr5:51390058-51391329). HEK 293 cell were plated in 24 well plates with seeding density of 1×10^5^ cells/well and incubated overnight. Each well was co-transfected using Lipofectamine 2000 (ThermoFisher, Cat. No. 11668019) with 500 ng reporter and 50 ng pRL Renilla Luciferase Control Reporter vector (Promega Cat. No. E2231). Luciferase intensity was measured 24 hours after transfection with Dual-Luciferase® Reporter Assay System (Promega, Cat. No. E1910) according to manufacture procedure in Orion L Microplate Luminometer with a 96 well plate (Corning Costar plate Cat. No. 3917). Firefly luciferase value were normalized for Renilla luciferase value. EZH2 was overexpressed using 100 ng of pCMVHA hEZH2 (Addgene Plasmid #24230) co-transfected with the plasmid of interest.

### Chromatin immunoprecipitation and qPCR (Chip-qPCR)

Chromatin immune precipitation was performed with Magna ChIP A/G immunoprecipitation kit (Merk, Catalog # 17-10085) according to manufacture procedure with little adjustment from the original protocol. In brief, 1.5×10^7^ HEK 293 cells were fixed in 1 % formaldehyde for 10 min at room temperature. Chromatin was sheared using a Covaris LE220 and sonication conditions were adjusted to reach chromatin fragment size between 200 and 500 base pare (bp). Shared chromatin was incubated overnight with anti EZH2 antibody (ThermoFisher, Cat. No. #49-1043) and control anti IgG antibody (Diagenode, Cat. No. #C15400001-15). Pulled-down DNA fragments were analyzed via qPCR using iTaq Universal SYBR Green Supermix (Bio-Rad, Cat. No. 1725121), primers are shown in Supplementary Information.

### EZH2 knock down

siPOOL targeting *EZH2* mRNAs (NM_001203247, NM_001203248, NM_001203249, NM_004456, NM_152998, XM_005249962, XM_005249963, XM_005249964, XM_011515883, XM_011515884, XM_011515885, XM_011515886, XM_011515887, XM_011515888, XM_011515889, XM_011515890, XM_011515891, XM_011515892, XM_011515893, XM_011515894) and control siPOOL were ordered at siTOOLs BIOTECH (Cat. No. 2146-EZH2). Reverse transfection in HEK 293 cells was performed with 3 nM final siRNA concentration with Lipofectamine™ RNAiMAX (ThermoFisher, Cat. No. 13778075) according to manufactural procedure. Cells were cultured in 6 wells plates (ThermoFisher, Cat. No. 140675) with seeding density of 0,2×10^6^ cells and were harvested for analysis after 48 hours of incubation.

### Western Blot

HEK293 cell protein was extracted with 0.3 mL of RIPA buffer (ThermoFisher, Cat. No. 89900) containing Protease Inhibitor Cocktail (ThermoFisher, Cat. No. 78410) from each well of a confluent 6 well plate according to manufactural procedure. The protein-lysate was measured with Pierce BCA Assay Kit (ThermoFisher, Cat. No. 23225). 30 µg of protein were diluted in Laemmli buffer and loaded in a 4 % to 15 % gradient mini-protean TGX gel (BioRad). After electrophoresis run, the protein was transferred on a PVDF membrane by Trans Turbo Blot (BioRad). The membrane was then blocked in 1 x TBS with 0.1 % Tween20 and 5 % milk powder solution for 2 hours at room-temperature, cut horizontally to separate EZH2 and beta-actin and then incubated at 4°C with primary EZH2 antibody (ThermoFisher, Cat. No. #49-1043; 1: 1.000) and anti-beta actin antibody (Sigma-Aldrich-A2228-RRID:AB_476697; 1: 50.000). Bands were visualized using enhanced chemiluminescence kit (SuperSignal West, Pierce).

### Genotyping of *ezh2^ul2^* larvae line

Larvae were derived from natural spawning of adults *ezh2^ul2+/-^*(Dupret et al., 2017) and were dechorionated at 48 hfp with 2 mg/ml pronase (Merk cat # P5147) in 0.3x Danieau for 10 min at room temperature (RT). Sub sequentially, 96 larvae were anesthetized with Tricaine and a fragment of caudal fin and tissue of each larva was surgically cut under a stereo microscope with a scalpel. DNA was extracted placing each dissection in 15 µL of 50mM NaOH solution and transferred in a 96 well plate (Star Lab cat. # 19103). In parallel, each cut larva was placed in a 96 well plate with 300 µL of Danieau water. To keep track of the genotype, the position where larvae are placed reflects the one of the plate with the tail tissues. Afterwards, the plate with tail and NaOH solution was heated at 90°C for 20 min, and cooled down to RT before adding 1/10^th^ volume of 1M Tris-HCl pH 8. PCR was performed with HOT FIREPol® Blend Master Mix Ready to Load (Solis Biodyne, Cat. Num. 04-25-00S20). Insertion of 22 nucleotides for hetertozygous (*ezh2^ul2+/-^*), homozygous (*ezh2^ul-/-^*) and WT was screened in a 2.8 % Phor Agarose (Biozym, Cat. No. 850180). Zfl are separated and collected for further analysis.

### Retro transcription quantitative PCR (RT-qPCR or qPCR)

For HEK293 cells, 1 mL of Trizol (ThermoFisher Cat. No. 15596026) was added in confluent siRNA and control culture (1.5*10^6^ cell/well) and RNA was extracted according to manufactural procedures. For zebrafish, 20 zfl were harvested after genotyping and RNA was extracted adding 1mL of Trizol. Whole or trunked larvae tissue was mechanically disrupted in the Trizol solution with Percellys 24 tissue homogenizer (peqlab) using 3 cycles of 10 seconds at 6.000 rpm. After that, RNA was extracted as manufactural procedures. Both for HEK293 and zebrafish larvae, cDNA was synthetized with 1µg of total RNA using iScript™ Reverse Transcription Supermix for qRT-PCR (Bio Rad, Cat. No. 1708841) according to manufactural procedures. For qPCR, final concentration 100 ng of cDNA was used for expression analysis with iTaq Universal SYBR Green Supermix (Bio-Rad, Cat. No. 1725121). qPCR was performed in a CFX96 Bio Rad reader with following cycles: 10 min of initial denaturation at 95°C followed by 40 amplification cycles consisting of 10 sec at 95°C and 1 min at 60°C. All data were normalized for beta acting reporter gene.

### In situ hybridization

In situ is performed and imaged as previously described in Zhang et al., 2017. Since different genotypes are used in the same in situ, to avoid change in staining due to probes pipetting error, fish were previously genotyped and 6 larvae (2 WT, 2 Ezh2+/- and 2 Ezh2 KO) were placed in the same probe and staining solutions.

### Immunohistochemistry

Zfl were fixed in 4 % paraformaldehyde PBS solution overnight, embedded in paraffin and cut in 3 µm thick slices. Staining was performed with the fully integrated staining solution Benchmark Ultra (Roche/Ventana). Slices were then incubated with anti-EGFP antibody (Takara Clontech 6323380; 1: 5.000) and anti ISL1 (AbCam cat no ab86472; 1: 1.000).

### Zebrafish larvae (zfl) imaging

Genotyped *Tg(wt1b:eGFP) ezh2^-/-^* and *Tg(wt1b:eGFP)* were incubated with 0.2 mM 1-phenyl 2-thiourea (PTU) Danieau solution to prevent pigmentation, anesthetized with 0.03 % tricaine (Merk) and mounted in 1.25 % low-melting temperature agarose. 3D images (both for live imaging and immunohistochemistry) were taken with a A1R HD25 ECLIPSE Ti2E laser scanning microscope using the NIS-Elements 5.21.02 software.

### Statistics

For HEK293-derived data, technical replicates per experiment were at least n = 3, and biological / independent experimental replicates were at least N = 3. For zebrafish-derived data, minimum 20 larvae (n ≥ 20) per condition were used for qPCR and independent biological replicates consists of N = 3. Statistical analysis was performed in GraphPad Prism 8.0. Luciferase assays and zebrafish qPCR were analyzed with ANOVA using as reference the parallel pGL3 control or the WT genotype respectively. T-test was used to compare qPCR of EZH2 siRNA with control and for ChIP-qPCR, where each EZH2 targeted region was compared with its IgG control. P-value is set as following: * ≤ 0.05; ** ≤ 0.01; *** ≤ 0.001 and **** ≤ 0.0001. Non-significant value (ns) corresponds at P-value > 0.05.

## Authors contribution

E. M. B.O. R.H. G.D. and H.R. designed the experiments; E.M. performed the experiments; E.M. B.O. and H.R. wrote the manuscript; N.H. and E.M. analyzed the human genomic data; T.F. Ö.Y. and T.L. performed assistant work for the experiments. P.-O. A. generated the zf Ezh2 knock out line. G.D. R.H. P.-O. A. B.O. and H.R. revised the manuscript.

## Author Approvals

All authors have seen and approved the manuscript. This manuscript and its data have not been accepted or published elsewhere.

## Acknowledgment

We are thankful for the support of the Zebrafish Core Facility (Bonn Medical Faculty). *Tg(wt1b:EGFP)* zf were provided by Prof. Christoph Englert (Leibniz-Institut, Jena). This study was in parts presented at the 2022 annual International Developmental Biologist conference (Algarve, Portugal).

## Data availability

The original contributions presented in the study are included in the article/Supplementary Materials, further inquiries can be directed to the corresponding authors.

## Ethics declaration

Animal husbandry and experimental setups were in accordance with European Legislation for the Protection of Animals used for Scientific Purposes (Directive 2010/62/EU). National law exempts all zebrafish experiments performed in larval stages up to five dpf before independent feeding from ethical approval.

## Supplementary information

**Supplementary Figure 1.**
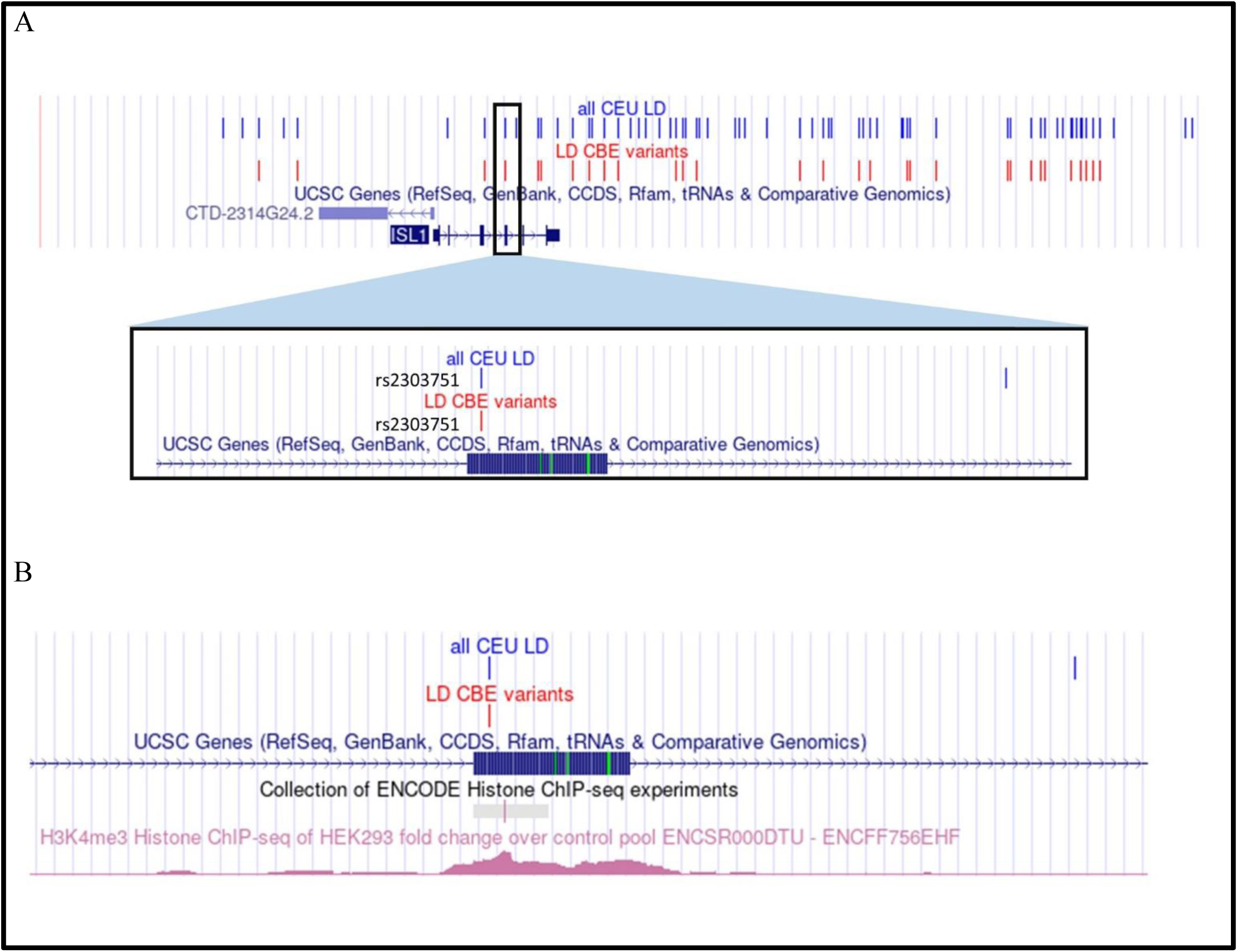
Map of the linkage disequilibrium variants and CBE associated variants in the ISL1 candidate region. A) Top panel: All CEU LD (in blue) shows all the variant in linkage disequilibrium (LD) with the top CBE associated rs6874700 variant; LD CBE variants (in red) shows CBE GWAS variants that are in common with the ones in LD. Bottom panel: zoom in the region where the only regulatory variant rs2303751 reseeds. B) H3K4me3 marker in the rs2303751 region displays a significant peak (Encode ID: ENCSR000DTU).

**Supplementary Figure 2.**
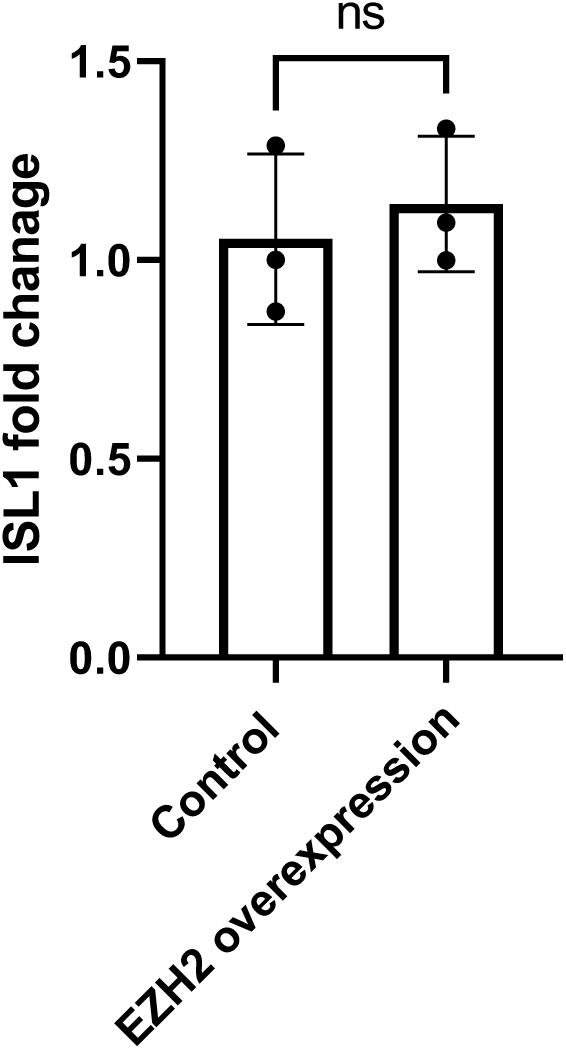
*ISL1* expression under EZH2 overexpression. qPCR against *ISL1* under EZH2 overexpression in HEK293 cells shows no difference in *ISL1* gene regulation compared to control.

### Oligonucleotides

**Table 1.**
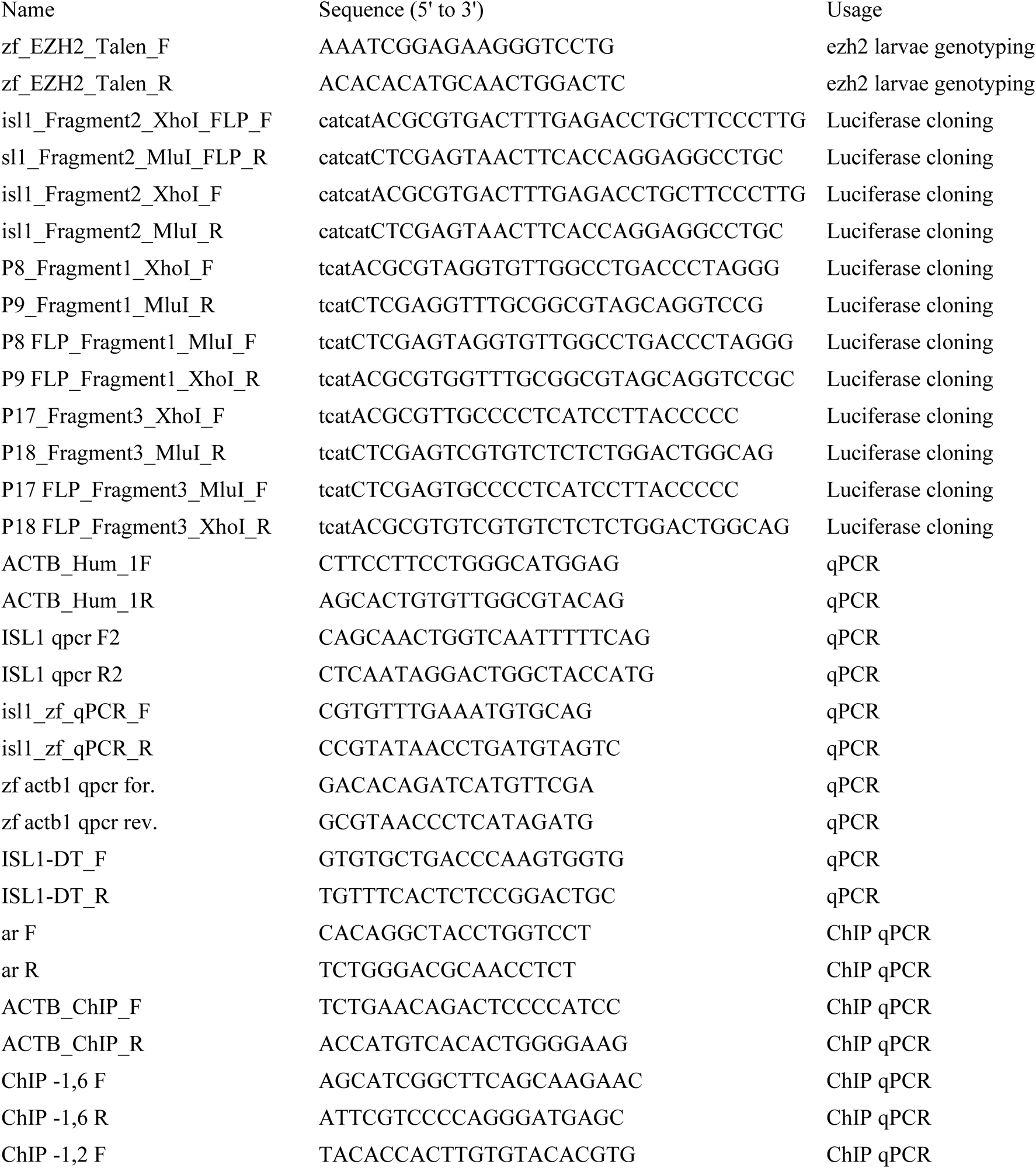
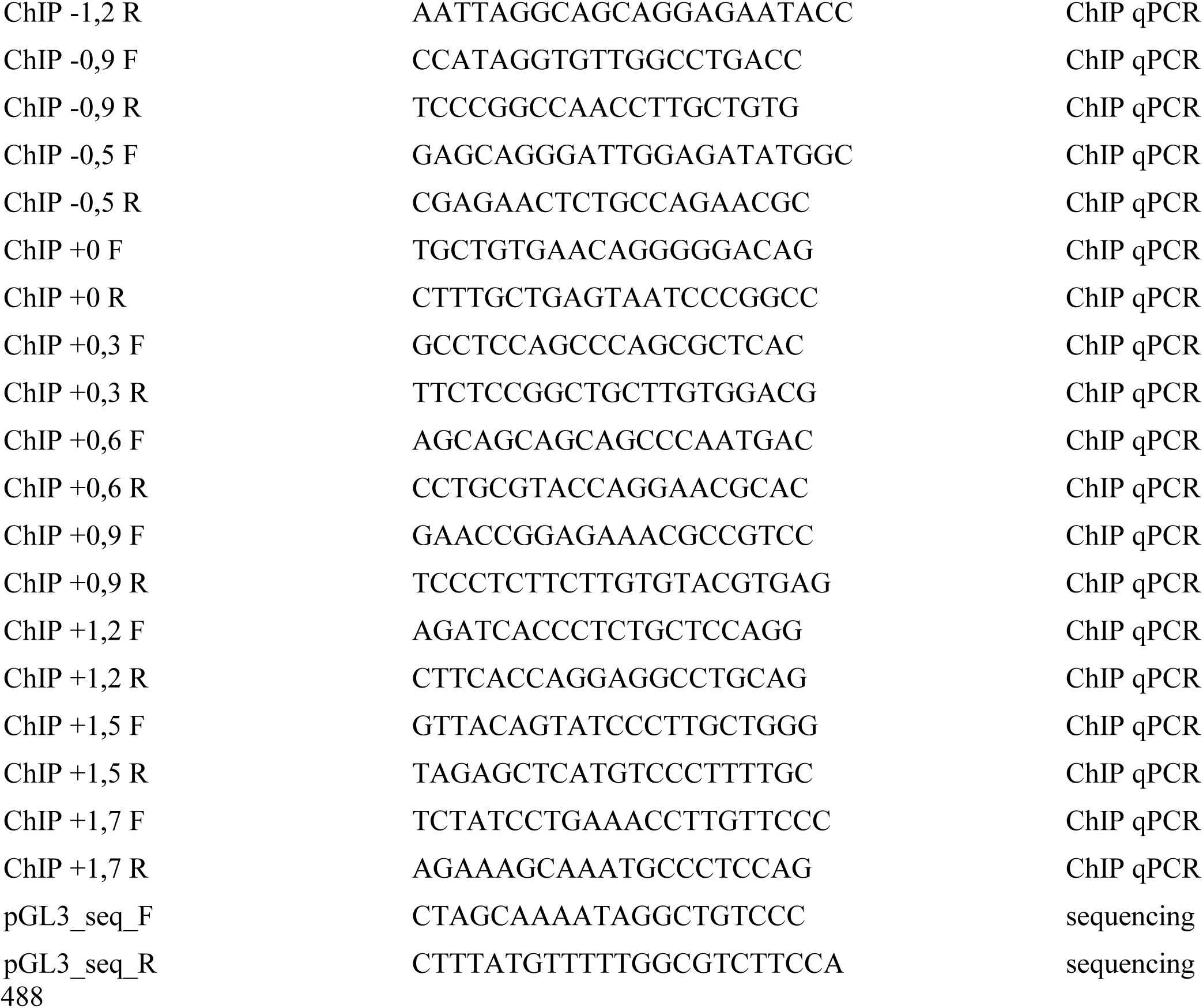

